# Advanced human cerebral organoids as a model for investigating glioma stem cell interactions with microglia and vascular cells and response to radiotherapy

**DOI:** 10.1101/2024.11.28.625826

**Authors:** Jérémy Raguin, Thierry Kortulewski, Oriane Bergiers, Christine Granotier-Beckers, Laure Chatrousse, Alexandra Benchoua, Laurent R. Gauthier, François D. Boussin, Marc-André Mouthon

## Abstract

The recent development of human brain organoids from induced pluripotent stem cells (IPSCs) enables the modeling of brain biology and pathophysiology, such as gliomas. However, most models lack vascular and/or immune systems, both of which play essential roles in maintaining brain health and in pathophysiological mechanisms. We have established a new method for generating vascularized complex cerebral organoids (CCOs) containing microglial cells (brain-resident macrophages) by incorporating bipotent hematopoietic/endothelial progenitors derived from the same IPSC lines during the early stages of development. This approach led to the formation of extensive vascular-like structures with blood-brain barrier characteristics, which were perfused upon transplantation into immunodeficient mice. Additionally, microglial cells exhibiting typical phenotypes and functionalities also developed within the CCOs. By coculturing CCOs with glioma stem cells, we demonstrated that this model effectively recapitulates the tumor niche of glioblastoma, showing vascular co-option, reprogramming of microglia into tumor-associated macrophages, and recurrence after radiotherapy. In conclusion, our vascularized and immunocompetent CCO model will be invaluable for understanding human brain development, exploring how this process is disrupted in diseases like gliomas, and discovering new therapeutic strategies.

## Introduction

Glioblastoma (GBM) is a brain tumor that is fatal despite aggressive treatment combining surgery, radiotherapy and chemotherapy. Glioma stem cells (GSCs), a subpopulation of tumor cells with stem cell properties are thought to play a major role in tumor relapse due to their resistance to treatment and invasive capacity [6, 47].

The innate immune and vascular systems within the tumor microenvironment also play a key role in GBM’s resistance to treatment [36] and undergo significant alterations following radiotherapy [21]. Indeed, tumor-associated macrophages (TAMs) represent the majority of non-tumor cells in glioblastoma (GBM), and their abundance correlates with disease severity, poor survival and recurrence [1, 18]. TAMs create immunosuppressive and pro-angiogenic conditions through the production of cytokines, promoting the proliferation and survival of GBM cells by synthesizing growth factors [36]. TAMs represent a heterogeneous population originating either from microglial cells (resident macrophages) or from circulating monocytes and/or bone marrow-derived cells [35]. The nature of TAMs depends, in part, on the genetic status of the tumor. In GBMs (isocitrate dehydrogenase (IDH) gene wild-type tumors), TAMs are predominantly monocyte-derived, whereas in IDH-mutated astrocytomas, they are more associated with microglial cells [2]. Vessel co-option and vascularization are among the strategies employed by some GBM cells to further invade the brain and regulate their proliferation and survival [38]. In the peri-necrotic regions of GBM, an abundance of destabilized microvessels polarizes macrophages towards a hypoxic state, which in turn increases vascular hyperpermeability, disrupting the efficacy of antitumor drugs [50].

Altogether these mechanisms contribute to the failure of various therapeutic approaches for GBM, but the exact roles of the GBM microenvironment in tumor development and relapse after treatments remain incompletely understood. Their elucidation requires study models that replicate its complexity in humans, which are currently not available.

The recent development of human brain organoids derived from human-induced pluripotent stem cells (IPSCs) allows for the modeling of brain biology and GBM pathophysiology [10]. Indeed, using 3D co-culture systems of human GSCs and brain organoids enables the identification of invasion signatures similar to those observed in surgical samples from GBM patients [23, 27]. However, 3D co-culture models combining GBM cells and brain organoids have certain limitations, particularly the absence of vascularization and immune cells. On one hand, vascularized organoids have been obtained using various sources of endothelial cells, genetically modified IPSCs [8], or after their transplantation into mice [29]. On the other hand, microglia-containing cerebral organoids have been generated (reviewed in [54]) with functional microglial cells [33] that closely resemble their in vivo counterparts [39]. Recently, cerebral organoids containing both vascular systems and microglial cells have also been created by assembling vascular organoids and cerebral organoids [43].

Brain vasculogenesis, the de novo formation of blood vessels, begins during embryoid development with the emergence of hemogenic endothelial cells (HECs) in the mammalian extraembryonic yolk sac [30]. Brain-resident macrophages, or microglia, also originate from this wave of HECs in the yolk sac and colonize the brain during early embryoid development [16]. The emergence of these HECs, which are of mesodermal origin, has been modeled using IPSCs [4]. Indeed, several research groups have developed strategies to derive HECs, inspired by human embryoid development [45]. HECs display a dual endothelial-hematopoietic phenotype characterized by the expression of VEGFR2/KDR/CD309 and the endothelial-specific junctional protein vascular endothelial cadherin (Cdh5/CD144) [5]. HECs also have the capacity to differentiate into various mesodermal cell types, including pericytes [5].

In this study, we developed a novel method to create vascularized cerebral organoids containing microglial cells from IPSCs derived from healthy donors, with the aim of establishing a new in vitro model for glioblastoma. Our strategy was to recapitulate the stages of embryoid brain development, including the colonization of cerebral organoids by HECs. By incorporating HECs during the early stages of cerebral organoid formation, we generated complex cerebral organoids containing both vascular structures and microglia. Finally, the invasion of GSCs within these vascularized, immunocompetent cerebral organoids as a model to recapitulate the GBM tumor niche.

## Material and Methods

### IPSCs culture

IPSC lines PDF01 (ISTEM, Evry, France) and GM25256*G (Coriell Cat#GM25256, RRID:CVCL_Y803) were cultured, respectively, in IPS-BREW XF (Miltenyi Biotec Cat#130-104-368, StemMACS™ iPS-Brew XF, human) on Vitronectin (Gibco^TM^ Cat#A14700, VTN-N) coated dishes and in mTESR1 (Stem Cell Technologies Cat#85850, mTeSR™1) on Matrigel (Corning Cat#354277, Corning® Matrigel® hESC-Qualified Matrix, LDEV-free) coated dishes. IPSCs were passaged and used at 70-80% of confluence. Pluripotency was routinely checked for >85% TRA1-81 (R and D Systems Cat# IC8495T, RRID:AB_3662726) and SSEA3 (R and D Systems Cat#FAB1434G, RRID:AB_3646484) staining by flow cytometry.

### Hemogenic endothelial cell differentiation

Hemogenic endothelial cells (HECs) were generated according to Vargas-Valderamma’s protocol with some modifications [45]. Briefly, IPSC colonies were moderately dissociated with 0.5mM EDTA (Invitrogen Cat#15575-038) and, optionally with Accutase^TM^ (Sigma Aldrich, Cat# A6964). One million of cells were cultured in 5mL STEMdiff™ APEL™2 medium (Stem Cell Technologies, Cat# 05275) in the presence of 10µM Y-27632 (Stem Cell Technologies Cat#7230), 50ng/mL Human VEGF-165 Recombinant Protein (PeproTech® Cat#100-20-050u) 50ng/mL Human BMP-4 Recombinant Protein (PeproTech® Cat#120-05-100u), 6µM CHIR99021 (BioTechne Cat#4423) in a 25 cm² flask. Alternatively, GM25256 cells for which dissociation is critical, HEC cultures were cultured into 24-wells AggreWell™800 (Stem Cell Technologies Cat#34815) with 500 cells/µwell. After 48h, cells were transferred to APEL2 medium with 50ng/mL VEGF-165, 50ng/mL BMP-4, 20ng/mL Human Flt-3 Ligand (FLT3L) Recombinant Protein (PeproTech® Cat#300-19-10u), 20ng/mL Human SCF Recombinant Protein (PeproTech® Cat#300-07-10u) and 20ng/mL Human TPO (Thrombopoietin) Recombinant Protein (PeproTech® Cat#300-18-10u) for 3 or 4 days. Five to seven days after their initiation, HECs were recovered from adherent clones by dissociation with TrypLE™ Express (Gibco Cat#12605010) or Accutase®.

### Obtaining of Complex Cerebral Organoids

Human cerebral organoids were generated according to Lancaster’s protocol with modifications [25]. Briefly, IPSC colonies were dissociated with 0.5mM EDTA and Accutase®. Embryoid bodies (EB) were formed with 9,000 cells in Corning® 96-well Clear Round Bottom Ultra-Low Attachment Microplate (Corning Cat#7007) and cultured in hESC medium (DMEM/F12 1% Glutamax [Gibco Cat#31331028] containing 20% [v/v] Knockout Serum Replacer [Gibco Cat#10828028], 1% [v/v] MEM-NEAA [Gibco Cat#11140050], 100μM 2β-mercaptoethanol [Gibco Cat# 31350010], 15% Fetal Bovine Serum [Gibco Cat#10500064], 1% [v/v] Anti-Anti [Gibco Cat#15240062]) with 10 µM Y-27632 and 4ng/mL thermostable human basic fibroblast growth factor (bFGF, Gibco Cat#PHG0360). The fifth day, EB were transferred in hESC medium without bFGF or Y-27632 for 6 hours. Then, EB were transferred in Neural Induction Medium (DMEM/F12 1% Glutamax containing 1% [v/v] N2 [Gibco Cat#17502048], 1% [v/v] MEM-NEAA, 1μg/mL Heparin [Stem Cell Technologies Cat#07980], 1% [v/v] Anti-Anti). Seven days after their initiation, embryoid bodies were cocultured for 2 days with dissociated HEC (10^4^ HEC/embryoid body) in neural induction medium. Then, complex cerebral embryoid bodies were embedded into 20µL droplet of GFR Matrigel (Corning Cat#354230) and cultivated into cerebral organoid differentiation medium (50% [v/v] DMEM/F12 with 50% [v/v] Neurobasal [Gibco Cat#21103049] containing 1% [v/v] Glutamax [Gibco Cat#35050061], 0.5% [v/v] N2, 2.63 µg/mL Insulin [Gibco Cat#], 0.5% [v/v] MEM-NEAA, 50µM 2β-mercaptoethanol) composed of 0.5% (v/v) B27 without vitamin A (Gibco Cat#12587010) supplemented with 20ng/mL VEGFA, 100ng/mL IL-34 (PeproTech® Cat#200-34), 10ng/mL GM-CSF (PeproTech® Cat#300-03). After 2-3 days, the medium was replaced by cerebral organoid differentiation medium with 0.5% (v/v) B27 (Gibco Cat#17504044), 20ng/mL VEGFA, 100ng/mL IL-34, 10ng/mL GM-CSF and renewed each 3-4 days. CCO were kept under orbital shaking from day 11 (85 rpm). After 17-18 days (i.e. 30-31 days after their initiation), CCO were maintained in cerebral organoid differentiation medium without growth factors and renewed each 3-4 days.

### Subcutaneous implantation of organoids into mice

All animal experiments described in this study were approved by the Institutional Animal Care & Use Committee of CEA/DRT (CEtEA) and were performed under the EU guidelines with approved protocol (APAFIS#40886-2023021010503692). Male NOD-SCID mice (RRID:IMSR_JAX:005557) were bred in our animal facility. Thirty days after their initiation, CCO were implanted into the leg of NOD-SCID mice. Mice were placed into a chamber by providing 2% isoflurane for anesthetization. Then, mice were immobilized under a laminar flow hood to cut a small incision at each back leg. CCOs were subcutaneously placed into the incision of the right and left back legs. Once CCO were inserted, the wound was closed with sutures and buprenorphine (0.1 mg/kg) was administered for pain relief. One month after xenotransplantation, mice were anesthetized and perfused with 25 mL of 70kDa FITC-dextran (Sigma Aldrich Cat#90718) for 5 min then euthanized. Subsequently, CCOs were recovered and fixed in 4% PFA for immunostaining.

### Flow cytometry

CCOs were dissociated into single cell according to Velasco’s protocol with modifications [46]. Briefly, CCOs were minced into small pieces by razor blade and incubated 30 min at 37°C, under orbital shaking (85 rpm), with 30U/mL Papain (Worthington Cat#LS003119), 100U/mL DNAse I (Sigma Aldrich Cat#D5025), 0.05mM EDTA (Sigma Aldrich Cat#E5134) and 0.18mg/mL L-cysteine (Sigma Aldrich Cat#C7352) in EBSS (Gibco Cat#24010043). After mechanical dissociation, cell suspensions were further incubated for 5-10 min at 37°C under orbital shaking. Then, cells were filtered into single cell in 10mg/mL Trypsin inhibitor type II (Sigma Aldrich Cat#T9128) and 100U/mL DNAse I (Sigma Cat#D5025).

Dissociated cells were incubated for 15 min at 4°C with fluorochrome-labeled antibodies (Tab. S1) in PBS 0.15% BSA. Then, cells were washed and resuspended in 0.15% BSA/PBS with 1µg/mL of bisBenzimide H 33258 (Sigma Aldrich Cat#23491-45-4 H) and sorted on a BD FACSAria II Cell Sorter (RRID:SCR_018934). Data were analyzed on FlowJo (RRID:SCR_008520).

### Glioma stem cell cultures

The TG16 and TG20 GSC lines were obtained from surgical resections carried out at Sainte Anne Hospital (Paris, France)[20, 40] on patients with high-grade IDH wt/mut gliomas according to the WHO classification (GBM: TG16; astrocytoma: TG20). GSCs were cultured as gliospheres in defined stem cell culture condition (serum-free Dulbecco’s Modified Eagle Medium DMEM/F12 supplemented with 1% Anti-Anti, 1x B27 without vitamin A, 5 μg/mL heparin, 20 ng/ml human recombinant epidermal growth factor (EGF, Sigma Aldrich Cat#E9644) and 20 ng/ml bFGF at 37 °C in an atmosphere containing 5% CO2. Every week, cells were dissociated after a 5 min incubation at room temperature with the Accutase cell dissociation reagent and reseeded at 0.5 × 10^6^ cells per T75 flask. GSC lines were transduced using a lentiviral vector containing a Luciferase/EGFP cassette (pTRIP-MND-Luciferase-Ires-GFP) [40].

### Coculture of GSC with CCO

Forty to 100 days after their initiation, CCOs were individually transferred into 1 well of 24-well plate (Falcon Cat#353047). Dissociated EGFP/Luciferase-expressing stable GSCs (2 x 10^4^ cells) were added to each well in 1mL of GSC medium without growth factor. Plates were incubated on orbital agitator (85 RPM). After 2 days, CCOs were washed in PBS and transferred into new 24 well conventional plate. Then, CCO/GSC were maintained under orbital shaking (85 RPM) in cerebral organoid differentiation medium without growth factor with biweekly renewal.

### Irradiation

Irradiations were performed with a GSR D1 irradiator (GSM) with four sources of Cs-137, with a total activity of around 180.28 TBq in March 2014, emitting 662 KeV gamma rays. Samples were irradiated at one single dose of 2 Gy (∼1 Gy/min) in 24-well plates. Prior to irradiation, dosimetry was performed in the same irradiation conditions with a cylindrical ionizing chamber 31010 by PTW as the recommendation of the AAPM’S TG-61. This ionizing chamber has a cavity of 0.125 cm3 calibrated in Cs-137 Kerma air with the PTB reference facility number 230451301. The polarity and the ion recombination were measured for this Cs-137 source. Each measurement was corrected using the KTP factor to account for variations in temperature and atmospheric pressure.

### Luciferase assay in CCO

Pre-warmed medium containing D-luciferin (PerkinElmer Cat#122799) was added to each well to reach a final concentration of 250 µg/mL and immediately processed for monitoring with a Bioluminescence Imager (Perkin Elmer IVIS LUMINA II). The flux (photons/sec) was measured each 2 min for 8 min, and the maximum value was recorded.

### Quantitative PCR (qPCR)

Cells were sorted into tubes containing RLT lysis buffer (Qiagen Cat#79216) and total RNAs were isolated with the RNeasy Micro Kit with DNase treatment (Qiagen Cat#74004) or alternatively with the NucleoSpin RNA XS (Marcherey-Nagel Cat#740902). For qRT-PCR experiments, total RNAs were reverse-transcribed into cDNA using the Reverse Transcription High Capacity Master Mix (Applied Biosystems Cat#4368814). q-PCR was performed on an Applied Biosystems StepOne Real Time PCR System (RRID:SCR_023455) using Power SYBR GREEN (Applied Biosystems Cat#4367659) with specific primers listed in Table S4. Data were analyzed on StepOne Software (RRID:SCR_014281).

### Manual and automated multiplex immunofluorescences

CCOs were fixed in 4% paraformaldehyde 3-4 hours at room temperature and conserved at 4°C in PBS with 0.2% azide. For 5µm slides, fixed CCOs were embedded in paraffin with Tissue-Tek (Sakura) and sliced with microtome (RM2125RT, Leica Microsystems), and were mounted on SuperFrost Plus (Epredia Cat#J1800AMNZ). After paraffin removal and citrate treatment, slices were blocked 1 h at room temperature in PBS with 10% FCS and 5% BSA (Sigma Aldrich Cat#A7906). The slides were incubated overnight with primary antibodies (Table S2) at 4°C. After PBS washing, the slides were incubated with secondary antibodies (1/1000, Table S3) at room temperature for 3 hours then washed with PBS. Alternatively, 4-plex was performed by automated immunostainnings on a Roche Ventana BenchMark ULTRA IHC/ISH System (RRID:SCR_025506) according to the manufacturer’s protocol (Zhang et al. 2017). Primary antibodies were the same as above and OmniMap anti-Ms HRP and anti-Rb HRP (Roche Cat#5269652001 and Cat#5269679001) secondary antibodies were coupled to OPAL520 (AKOYA Biosciences Cat# FP1487001KT), OPAL570 (AKOYA Biosciences Cat#FP1488001KT), OPAL620 (AKOYA Biosciences Cat#FP1495001KT), OPAL650 (AKOYA Biosciences Cat#FP1496001KT) reagents with Tyramide Signal Amplification. Slides were mounted with Fluoromount-DAPI (Southern Biotechnologies Cat#0100-20). Images were captured on a Leica SP8 LIGHTNING confocal microscope (RRID:SCR_018169) with the following objectives: ×20/0.75 NA, and ×63/1.3 NA Oil. Images were analyzed on Leica Application Suite X (RRID:SCR_013673) or IMARIS (RRID:SCR_007370) softwares.

## 3D Immunofluorescence and CE3D clearing

Fixed CCOs were embedded in 2% low melting point agarose in PBS and sliced at 500µm with a Leica VT1200S vibrating microtome (RRID:SCR_020243). Slices were stained and cleared following the Ce3D protocol with modifications [26]. Briefly, slices were blocked 24h with Alternative Blocking Buffer (ABB) (PBS-TABS [Sigma Aldrich Cat#18912] with 0,3% Triton X-100 [Sigma Aldrich Cat#T8787], 1% BSA, 1X BD Perm/Wash Buffer [BD Cat#554723], alternatively 1% of Donkey Serum [Eurobio Scientific Cat#CAEMTN00-0U] was added for secondary antibodies). Slices were incubated 72h with primary antibodies (Table S2) in ABB at 37°C under 85 RPM. After a 12 h washing with Conventional Washing Buffer (CWB) (PBS-TABS with 0.3% Triton X-100, 0.5% 1-thioglycerol [Sigma Aldrich Cat#M1753], 0.1% Saponine [Sigma Aldrich Cat#S-4521]) at 37°C under 85 RPM, slices were incubated 72h with secondary antibodies (1/1000, Table S3) and with 5µg/mL of DAPI (Sigma Aldrich Cat#D9542) in ABB at 37°C under 85 RPM. After a washing with CWB, slices were placed into 24 well plate glass bottom (Azenta Cat#4TI-0243), and were progressively cleared under orbital shaking (100 RPM) with 30%, 50%, 70%, 90% gradient of Ce3D solution in PBS, each for 1h30 then with 100% Ce3D solution overnight. Supernatant was eliminated and cover slips (Epredia Cat#CB00120RA120MNZ0) were placed. Images were acquired on a Spinning-Disk confocal microscope (Nikon, GATTACA) with a 1mm piezoelectric (Piezoconcept) with the following objectives: ×10/0.45 NA, ×20/0.75 NA, ×40/0.45 NA, and ×60/1.4 NA oil. Images were analyzed with IMARIS software.

### Data analyses

All data points were plotted, and the median along with the interquartile range (1st and 3rd quartiles) are shown. Statistical significance was determined using Kruskall-Wallis and uncorrected Dunn’s tests for multiple comparisons (P<0.05: *; P<0.01: **) or with Mann-Whitney (GraphPad Prism 9 [RRID:SCR_002798]) for pair comparisons (P<0.05: #; P<0.01: ##). For flow cytometry analyses, some CCOs had a low CD45-positive cell content (less than 0.040%) and/or contained less than 17% P2RY12-positive cells within the CD45+ population; they were therefore removed from the analyses.

## Results

### Integration of HECs did not hinder cerebral organoid development

We designed a stepwise protocol by separately inducing mesodermal differentiation into hemogenic endothelium and neuroectodermal differentiation from the same iPSC lines, then combining them to generate complex cerebral organoids (CCOs) (Figure 1A). HECs were obtained using a recently developed protocol with some modifications [45]. Hematopoietic-endothelial differentiation of iPSCs was confirmed by the expression of markers by FACS and RT-qPCR for CD309/KDR and CD144/Cdh5, and by FACS for CD31 and CD146, a mesenchymal/endothelial marker (Fig. SII1). RUNX1, a transcription factor involved in hematopoiesis, was also expressed at the mRNA level (Fig. SI1).

**Fig. 1:**
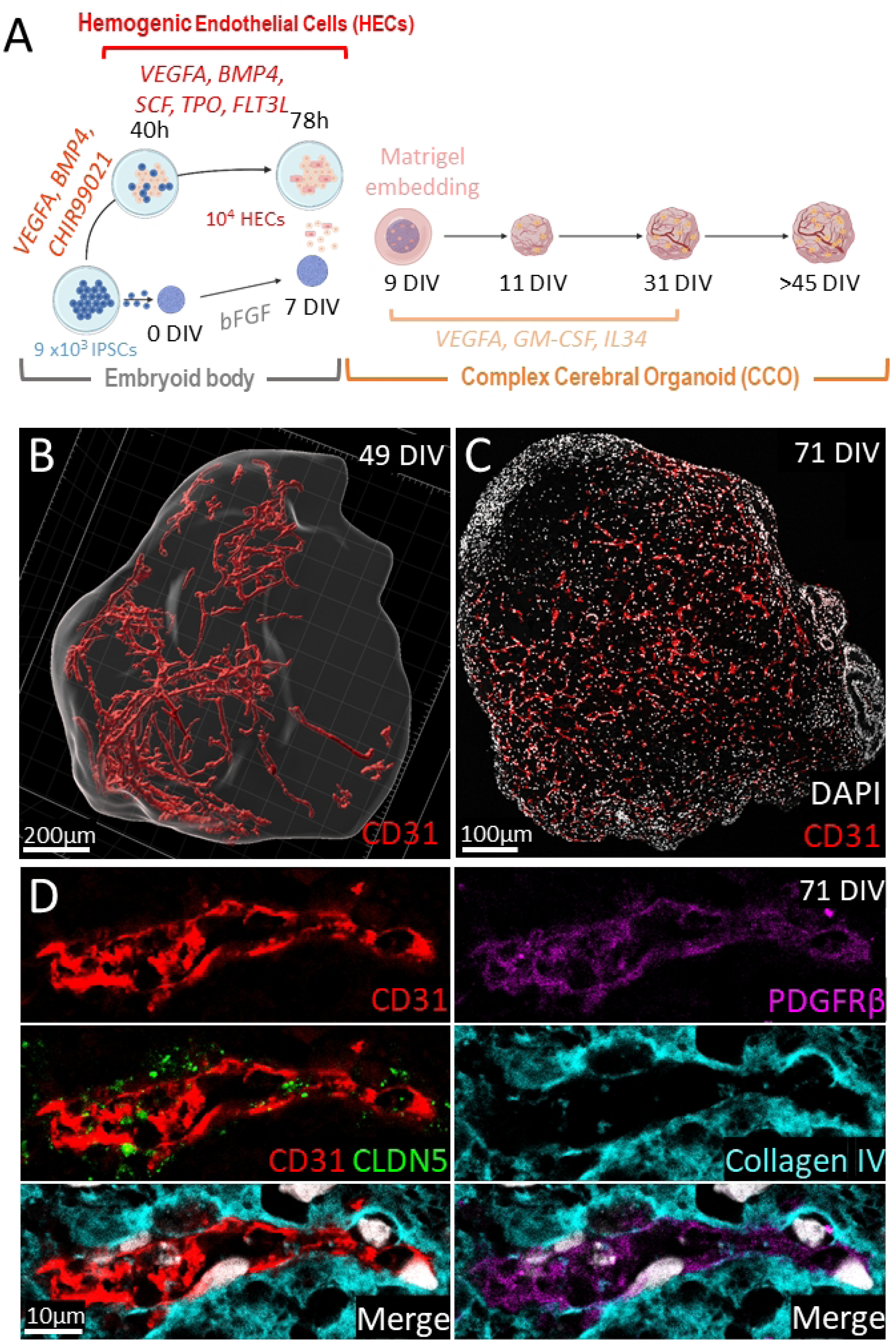
Vascular-like structures develop in CCO. (A) Schematic representation of the protocol to obtain CCOs. The same IPSC lines were used to generate hemogenic endothelial cells (HECs) and embryonic bodies. Then they were combined the 9^th^ Day In Vitro (DIV) after initiation of embryonic bodies. Immunostaining for CD31 were performed on 500µm sections (PDF01 line, a 3D modelling is shown in B) and on 5 µm sections (GM25256 line, C). (D) Multiplex immunostaining for CD31, PDGFRβ, CLDN5, Collagen IV

Meanwhile, embryoid bodies were formed from the same iPSC lines to derive cerebral organoids [25], which were then cocultured with HECs on day seven in the presence of VEGFA, IL-34, and GM-CSF for three weeks to support differentiation into vascular and microglial cells (Figure 1A). Since we showed that the CCOs intrinsically produced VEGFA, IL-34, addition of exogenous growth factors was thus discontinued at day 30 (Fig. SI2). Moreover, RT-PCR also suggest an endogenous production of CX3CL1 in the CCOs, which is involved in microglial homeostasis (Fig. SI2).

Given the presence of lineage-directing growth factors and the incorporation of HECs into the cerebral organoids, we examined whether neuroectodermal differentiation remained unaffected. CCOs were collected between days 50 and 110 after initiation and processed for immunostaining, either on thin slices or clarified thick slices, followed by 3D confocal microscopy. The CCOs contained neural rosettes composed of SOX2-positive neural progenitors and SOX9-positive gliogenic progenitors (Fig. SI3A-B). Neural precursors expressing doublecortin were abundant and widely distributed within the CCOs (Fig. SI3C). Differentiation into neurons was confirmed by the presence of β3-tubulin and MAP-2 immunostaining, which were widely present throughout the CCOs (Fig. SI3C). The presence of numerous astrocytes was confirmed by S100β/GFAP double-positive cells (Fig. SI3E), further corroborated by their co-expression with SOX9 (Fig. SI3F).

Altogether, these findings show that the incorporation of HECs did not hinder either neuronal or glial differentiation within the cerebral organoids.

### Pseudo brain vascularization in complex cerebral organoids (CCOs)

Fifty days after initiation—20 days after the removal of growth factors (VEGF, IL-34, GM-CSF)— numerous tubule-like structures positive for CD31, a specific endothelial marker, had extensively developed within the CCOs (Figure 1B-C). These structures occupied a significant portion of the organoids, including the center. These vessel-like structures were observed in CCOs initiated from all the iPSC lines tested and remained prominent at later time points (up to 70 days). Additionally, immunostaining for collagen IV was detected adjacent to CD31, resembling the basal lamina of blood vessels (Figure 1D).

To determine whether these vessel-like structures mimic the blood-brain barrier, we performed multiplex immunohistological analyses on thin CCO slices. Claudin-5 (CLDN5) and ZO-1, markers of tight junctions in brain vessels, were present in the CCOs in proximity to CD31 immunolabeling (Figure 1D and Fig. SI4). Furthermore, PDGFRβ positivity indicated the presence of pericytes surrounding the endothelial cells (Figure 1D).

To further examine the potential functionality of these vessel-like structures, CCOs were transplanted into immunodeficient NOD-SCID mice. One month post-transplantation, Dextran-FITC was infused intracardially into the mice’s vascular system, after which the CCOs were recovered (Fig. SI5A). IPSC-derived vessel-like structures in CCOs were identified using a specific anti-human CD31 antibody (Fig. SI5B). Figure S5C demonstrated that Dextran-FITC flowed through these vascular structures, indicating their connection to the mouse vasculature and functionality. These vessel structures were also covered by PDGFRβ-positive pericytes (Fig. SI5D).

Overall, our data show that the vascular-like structures in CCOs acquired blood-brain barrier features.

### Functional Microglia-like cells developed into complex cerebral organoids

Next, we analyzed whether the early incorporation of HECs into embryoid bodies effectively led to the generation of microglial cells in the CCOs. One month after initiation, numerous Iba1-positive cells, a pan-macrophage/microglia marker, were widely observed, extensively infiltrating the CCOs (Figure 2A). Virtually all Iba1-positive cells were also positive for P2RY12, a specific microglial marker (Figure 2B).

**Fig. 2:**
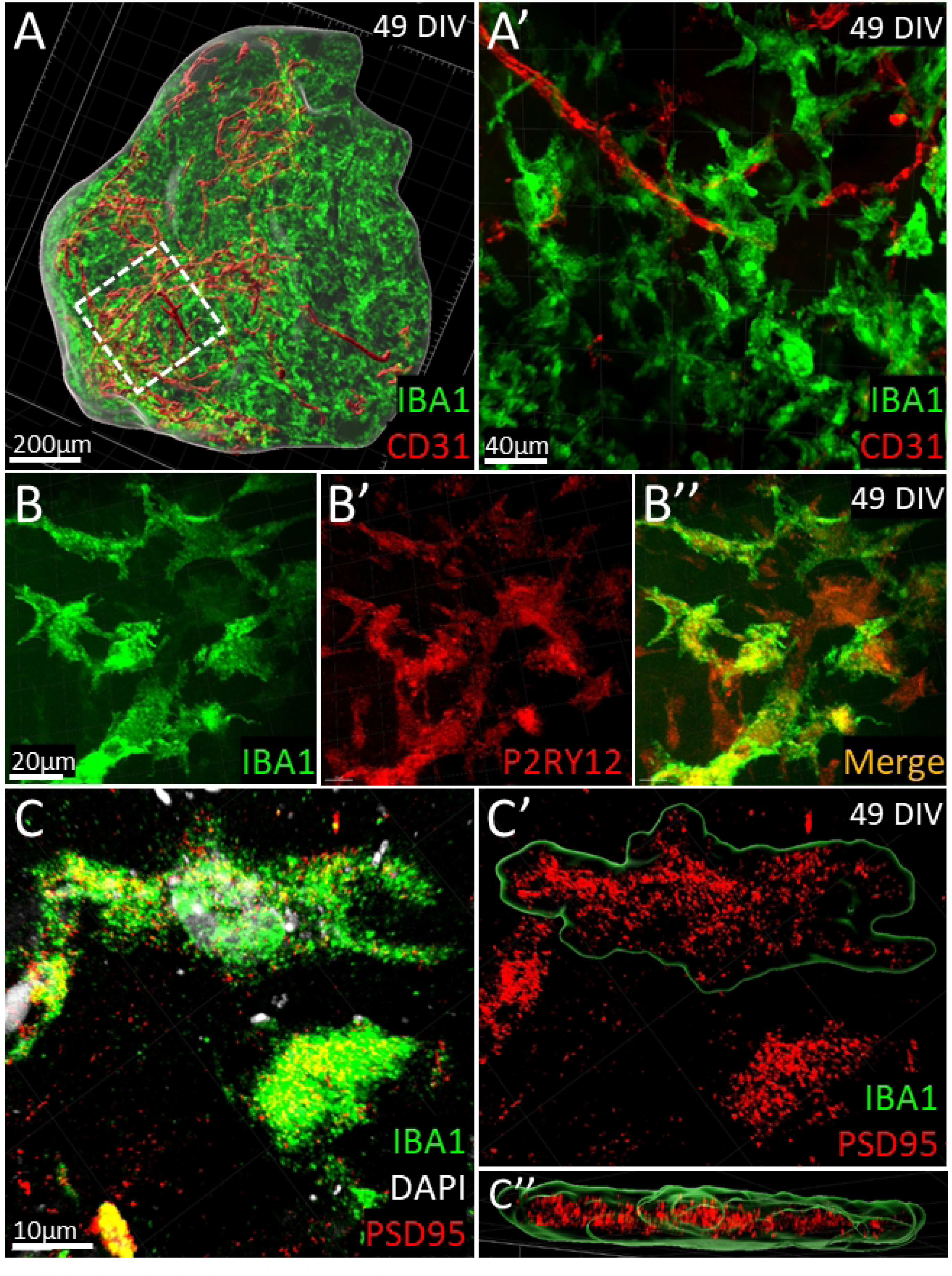
Microglial-like cells develop in CCO and exert a pruning feature. (A) Representation of immunostaining for CD31 and IBA1 on 500 µm section of the CCO (enlargement is shown in A’). (B, B’, B’’) Almost all Iba1-positive cells co-expressed the microglial marker P2RY12. (C, C’, C’’) Iba1 expressing cells showed internalization of the neuronal PSD95. The outline of a positive Iba1 cell is shown in C’ and C’’

To further characterize these microglia-like cells, we conducted FACS analysis of various microglial markers in single-cell suspensions obtained through enzymatic dissociation of the CCOs. The total number of cells per CCO peaked around 10 weeks after initiation, followed by a decline at later time points (Figure 3). While very few, if any, CD45-positive cells were detected in conventional organoids, significant levels of CD45+ cells were found in CCOs, with almost all being positive for CD11b, P2RY12, and CX3CR1 (Figure 3 and Fig. SI6). The number of CD45+CD11b+P2RY12+ cells peaked around ten weeks and subsequently decreased (Figure 3).

**Fig. 3:**
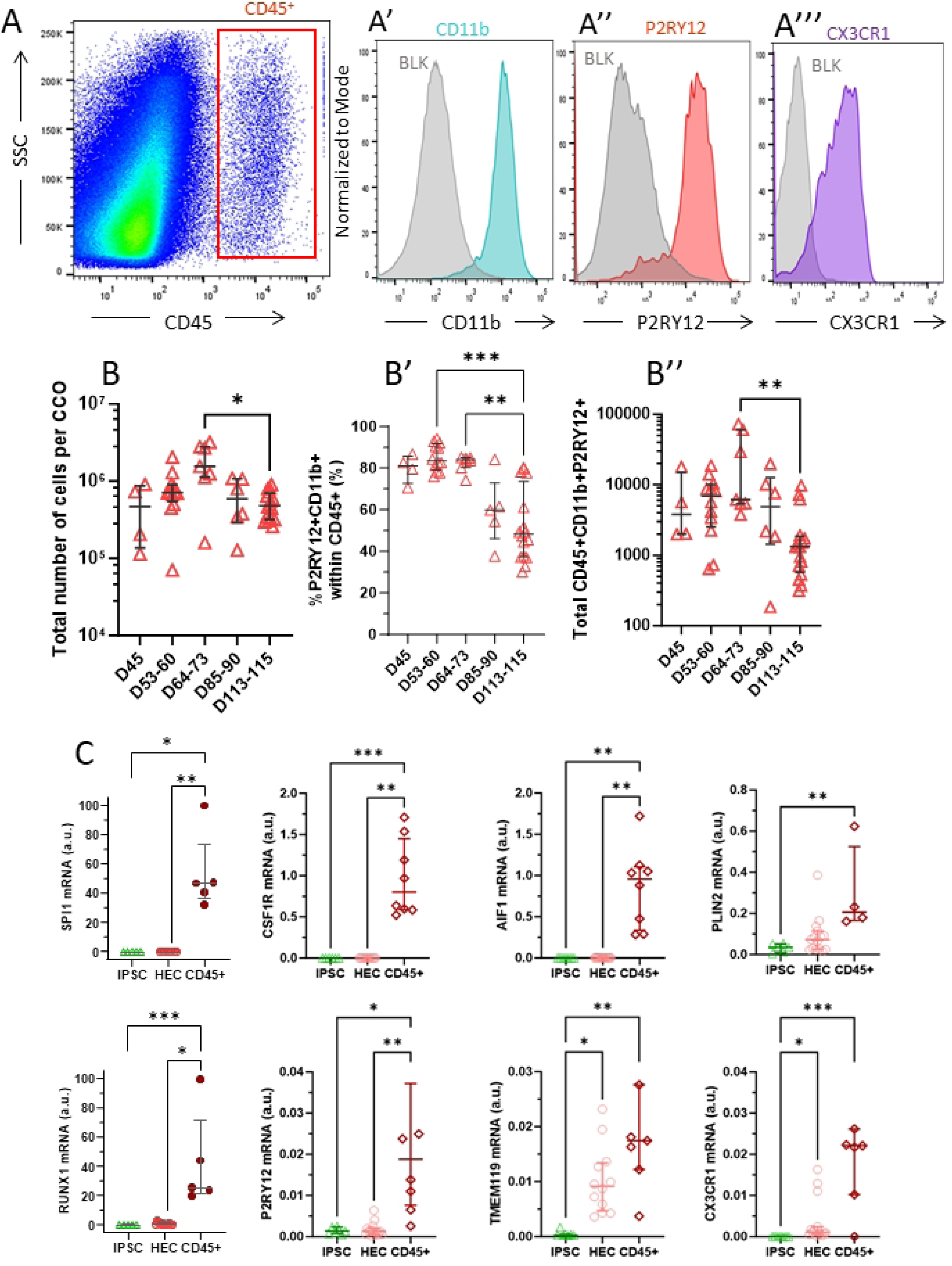
Flow cytometry analysis of microglial-like cells. (A) Microglial-like cells were contained in the CD45+ population expressing CD11b (A’), P2RY12 (A’’) and CX3CR1 (A’’’). (B) The amount of total cells per CCO was decreased at later time point. (B’) The great majority of CD45+ also expressed CD11b and P2RY12. (B’’) The total amount of iMG cells decreased at later time point. (C) Expression of microglial genes was determined by RT-qPCR in sorted CD45+CD11b+ iMG and compared to parental IPSC and to HECs (mRNA expression was normalized to 18S, a.u.: arbitrary unit)

We further examined the expression of macrophage/microglial genes in sorted CD45+ cells. The hematopoietic transcription factors SPI1 and RUNX1 were highly expressed in CD45+ cells compared to parental iPSCs and differentiated HECs (Figure 3C). Additionally, CSF1R and AIF1 were also enriched in CD45+ cells (Figure 3C). Other markers, including P2RY12, TMEM119, PLIN2, and CX3CR1, showed high enrichment in CD45+ cells compared to iPSCs and differentiated HECs (Figure 3C). Therefore, these CD45+/Iba1+ cells are referred to as induced microglia thereafter (iMG).

Microglial cells play an important role in synapse pruning and promoting brain maturation. To demonstrate that iMG/Iba1+ cells were functional, we first examined their ability to engulf synapses by double-staining for postsynaptic density protein 95 (PSD95) and Iba1. Remarkably, iMG exhibited numerous PSD95 puncta distributed throughout their cytoplasm, indicating synaptic engulfment (Figure 2C).

Overall, our data showed that the CCOs contained functional iMG.

### Coculture of GSCs with CCOs recreated a complex tumor niche

We next investigated whether our organoid model could effectively replicate the glioblastoma (GBM) tumor niche. GBM tumor niches consist of tumor cells, including GSCs, as well as non-neoplastic cells such as neural and vascular cells, and tumor-associated macrophages (TAMs). To this end, we performed 3D cocultures of CCOs with GSC lines derived from adult patients with wild-type IDH GBM (TG16) or IDH-mutated grade IV astrocytoma (TG20). Both GSC lines exhibited high expression by FACS of the stem cell marker SOX2, with TG16 showing elevated levels of CD44 (Fig. SI7), consistent with its mesenchymal phenotype [14]. Additionally, RT-qPCR showed that TG16 GSCs expressed higher levels of chemokines, cytokines, and immune checkpoints compared to TG20 GSCs (Fig. SI7). Following xenotransplantation into nude mice, both TG16 and TG20 GSC lines formed tumors in vivo characterized by vascular aberrations and a high infiltration of Iba1+ macrophages/microglial cells (Fig. SI8A-B). FACS analyses of dissected striatum showed a higher abundance of CD45^++^CD11b^+^ cells, corresponding to monocyte-derived TAMs, in tumor-bearing brains compared to controls (Fig. SI8C). Additionally, sorted CD45^+^ cells, encompassing both macrophages and microglia, exhibited elevated expression of key genes (MSR1, SPP1, TGFβ1), indicating their transition to an immunosuppressive TAM phenotype (Fig. SI8D) consistently with their expression of cytokines and immunomodulators (Fig. SI7).

GSCs were cocultured with CCOs (45-60 DIV) for two days under orbital shaking to enrich for cells demonstrating active adhesion. Two weeks after coculture, both TG16 and TG20 GSCs formed focal clones that were dispersed within the CCOs but were largely excluded from the densely packed neural rosettes (Figure 4A-B). Organoids were dissociated between 14 and 45 days after initiation to estimate the amount of GSCs by FACS. We observed a sharp increase in GSC content for the TG16 line, which was less pronounced for the TG20 line (Figure 4C).

**Fig. 4:**
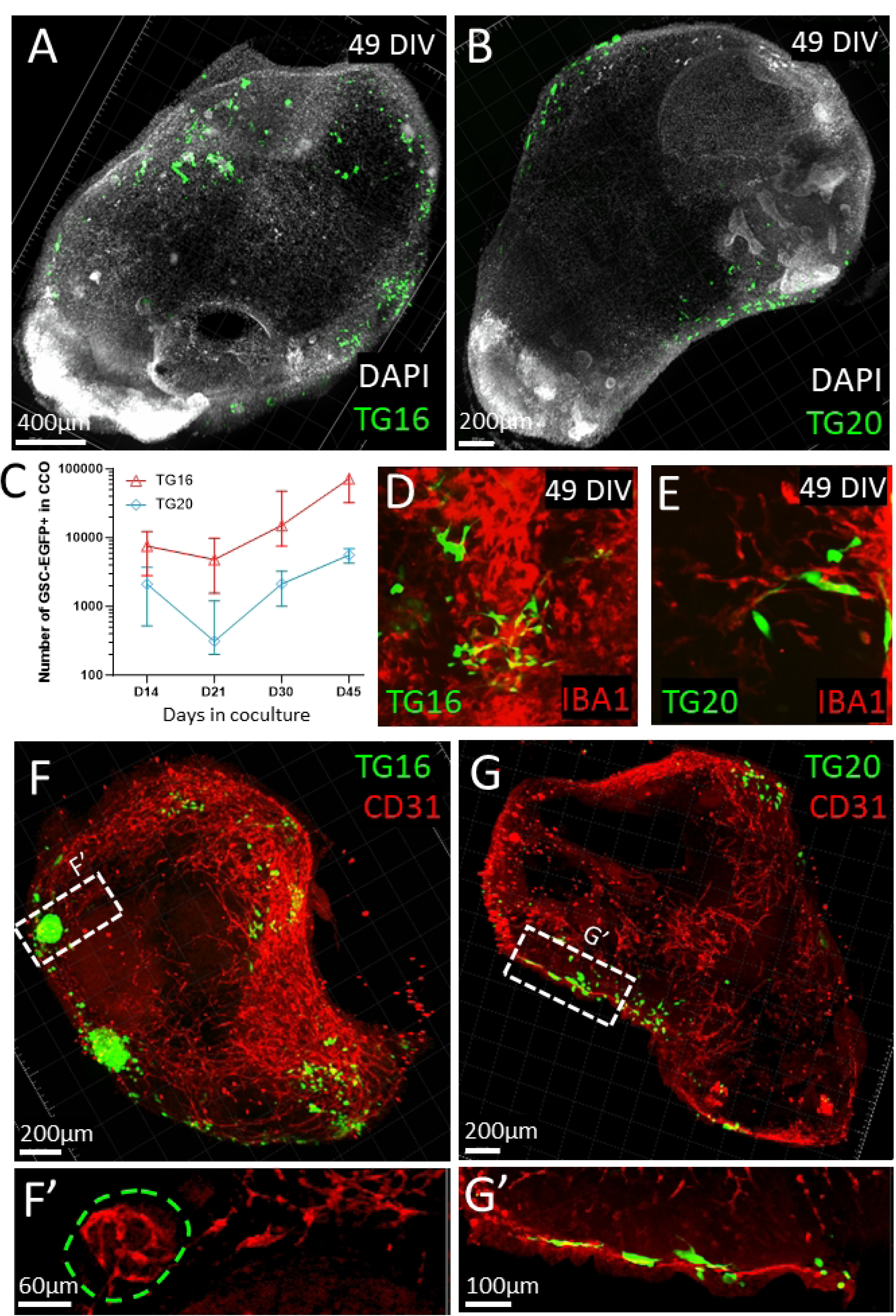
Cocultures of GSCs with CCO recapitulate the GBM tumor niche. (A) TG16-GSCs and (B) TG20-GSCs were cocultured with CCO for 4 weeks and were immunodetected for EGFP on 500 µm slices. (C) CCO were dissociated at different times after coculture and EGFP-GSCs were quantified by FACS. (D-E) GSCs were observed at the proximity of Iba1+ cells. (F-G) GSCs were in close contact to vascular-like CD31+ structures (enlargement in F’ and G’). (F’) The location of the TG16 clone is marked out

Resident microglial cells, which form part of the TAM population, along with vascular cells, play a critical role in GBM development and treatment resistance [36]. We observed that GSCs were located in close proximity to iMG in the CCO cocultures (Figure 4D-E). Additionally, GSCs were found in close contact with endothelial cells, showing a vascular reorganization, probably associated with vascularization within TG16 tumors (Figure 4F-F’), and evidence of vessel co-option in TG20 tumors (Figure 4G-G’).

Further dissociation of the cocultures enabled us to quantify the effects of GSCs on endothelial cells and iMGs by FACS. The percentages of CD31+ endothelial cells and iMGs were not significantly altered by the presence of GSCs compared to CCO alone (Fig. SI9). However, GSCs increased CD31 expression on endothelial cells (Fig. SI9), consistent with observations in the GBM tumor core [53]. Moreover, iMGs displayed an immunosuppressive phenotype in the presence of GSCs, particularly in the case of TG16. This was demonstrated by increased CD206 expression in FACS analyses and upregulated MSR1 mRNA levels (Fig. SI9). This shift was associated with higher expression of SPP1, a TAM marker (Fig. SI9).

In summary, these results demonstrate that cocultures of GSCs with CCOs effectively recreate key features of the GBM tumor niche.

### The complexity of CCO niche promotes GSC aggressiveness after irradiation

Radiotherapy is usually given as fractionated doses of about 2 Gy, a dose GSCs survive and exhibit diffuse invasion into neighboring brain tissues. Thus, we analyzed the effects of a single 2 Gy irradiation on CCOs two weeks after coculture with GSCs. The 2 Gy exposure resulted in a modest reduction in total cell numbers two weeks post-irradiation (Fig. SI10). To assess the effects of 2 Gy irradiation on GSC growth, we monitored luciferase bioluminescence, which correlated with the number of EGFP-labeled GSCs (Fig. SI11). We measured the bioluminescence of TG16 and TG20 GSCs in CCOs on days 16 and 23 after coculture, corresponding to days 2 to 9 post-irradiation, and compared it to conventional COs (Figure 5). The bioluminescence signal for TG16 and TG20 cells increased similarly in CCOs and conventional COs, indicating that GSCs developed comparably in each environment (Figure 5B and Fig. SI12). However, irradiation favored TG16-GSC proliferation in CCOs, while it had no effect on TG16 in coculture with conventional COs (Figure 5B). In contrast, irradiation tended to reduce TG20 growth in either conventional COs or CCOs (Fig. SI12).

**Fig. 5:**
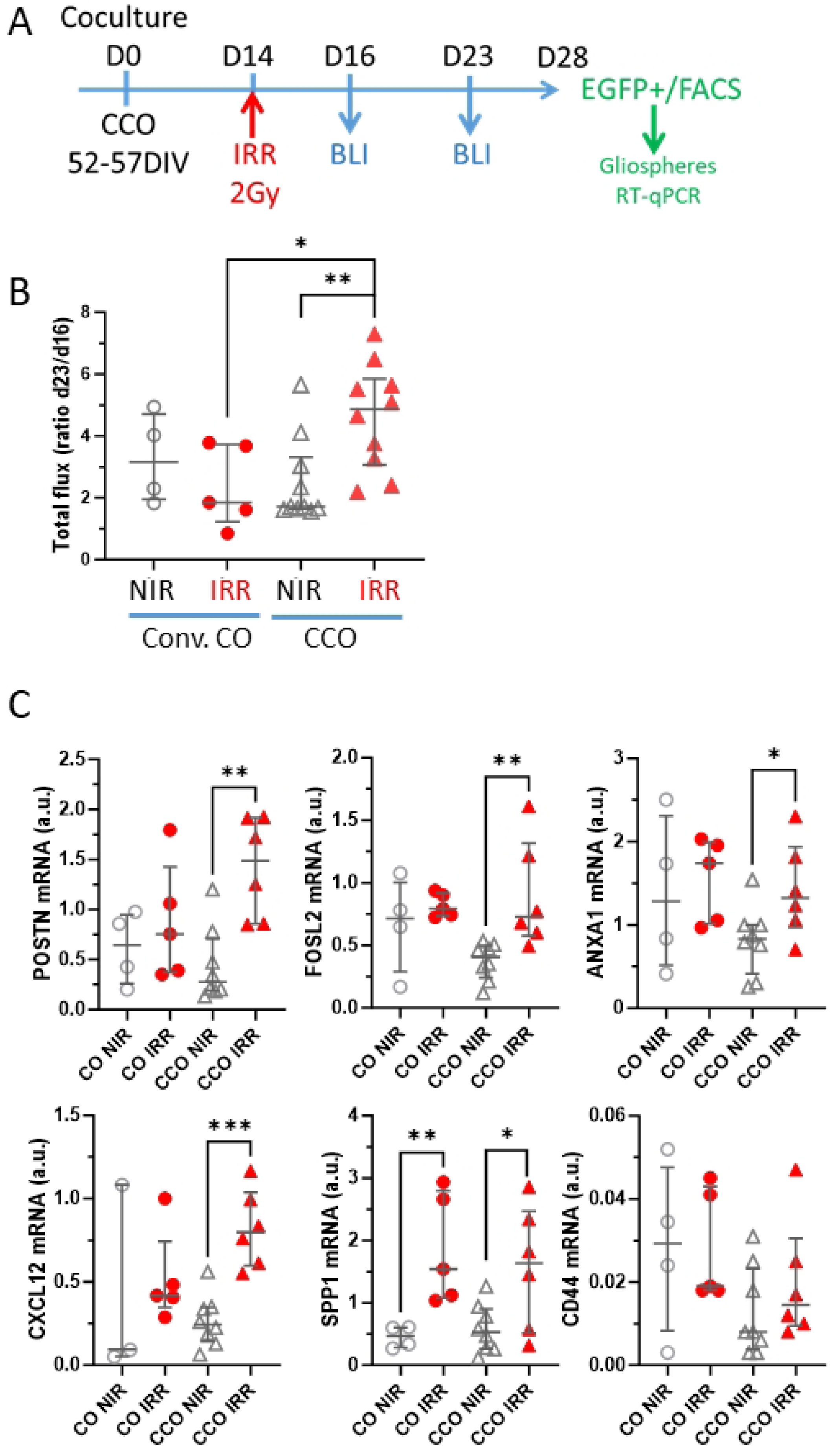
Irradiation promotes aggressiveness of GSCs in CCO. (A) TG16-GSCs were cocultured with CCO for 14 days then were irradiated (IRR), or not (NIR). The amount of GSCs was determined by Luciferase bioluminescence (photons/sec) 2 and 9 days later (D16 and D23). Then cocultures were dissociated and EGFP-GSCs sorted by FACS. (B) Longitudinal analyses of bioluminescence for GSCs cocultured with conventional CO (Conv. CO) or complex CCO. (C) GSCs were sorted and the gliosphere capacity was determined. (D) EGFP-GSCs were sorted and the expression of a set of genes was determined by RT-qPCR and compared to parental TG16 GSCs

We then analyzed the expression in sorted TG16 GSCs of a set of genes associated with GBM development and aggressiveness, known as the Natural Evolution Signature (NES) [52]. While CD44 mRNA levels were minimally impacted by irradiation, a specific set of genes (SPP1, POSTN, FOSL2, ANXA1, and CXCL12) was upregulated in TG16 cells cocultured with CCOs following irradiation (Figure 5C). In contrast, with the exception of SPP1, the expression of these genes remained largely unchanged in TG16 tumors derived from conventional COs (Figure 5C). This gene alteration aligned with the observed radiation-induced proliferation of TG16 GSCs.

We then analyzed the effects of irradiation on iMGs/TAMs using FACS. While the proportion of iMGs remained unchanged in CCOs (Fig. SI10), the expression of HLA-DR, CD11b, CD163, and CD206 on iMGs increased following irradiation, indicating their alternative activation toward an immunosuppressive TAM phenotype (Fig. SI10). This response to radiation was prominent in the presence of TG16 but minimal with TG20 (Fig. SI10).

## Discussion

To generate CCOs that contained both vascular structures and microglia-like cells, we replicated the stages of embryoid brain development, focusing on the colonization of cerebral organoids by endothelial cells and primitive macrophages. We thus incorporated bipotent hemato/endothelial cells (HECs) during the early stages of cerebral organoid formation. We validated these vascularized, immunocompetent cerebral organoids to model the GBM tumor niches and assess treatment effects. Given the critical role of vascularization in tissue physiopathology, recent efforts have focused on vascularizing cerebral organoids. Brain vasculature is formed by a combination of endothelial cells, astrocytes, pericytes, and neurons, creating the blood-brain barrier (BBB), an impermeable structure [12]. Most BBB cell types, such as astrocytes, pericytes, and neurons, are derived from neuroepithelium and can arise spontaneously within cerebral organoids. However, conventional cerebral organoids lack the ability to generate vascular endothelial cells, as these cells originate from the mesoderm.

In this study, we added iPSC-derived HECs along with exogenous VEGF and BMP4 to initiate endothelial differentiation. We anticipated that neural progenitor cells within the organoids would produce signals necessary for inducing vascular network formation [9]. Indeed, we observed vascular-like structures, identified by the endothelial marker CD31, surrounded by basal lamina (collagen IV) and pericyte coverage (PDGFRβ). Additionally, the presence of CLDN5 and ZO-1, key components of the tight junctions associated with the blood-brain barrier (BBB), was confirmed.

Recent studies have also modeled vascularized brain organoids with BBB features by fusing vascular and cerebral organoids in vitro [13, 43]. However, these vascular structures initially lacked a lumen, leading to apoptosis in the absence of blood flow. Consistent with previous findings [8], once our CCOs were transplanted into immunodeficient mice, the vascular structures became perfused with blood, confirming their functionality. In a previous study, iPSCs expressing ETV2 contributed to the formation of a complex vascular network within CCOs and enhanced the functional maturation of the organoids [8]. In our model, the inclusion of HECs enabled the generation of a substantial number of neurons within the CCOs. Moreover, we observed significant numbers of astrocytes (S100β/GFAP) as early as days 60-80, whereas astrocyte abundance is typically reported to increase after 100 days [34, 37]. This suggests that the combination of vascularization and microglia supports the maturation of cerebral organoids.

Microglial cells play a critical role in regulating neuronal wiring, angiogenesis, and immune responses in the brain, originating from extraembryonic erythroid-myeloid progenitors [44]. Various techniques have been used to generate cerebral organoids containing microglial cells [54]. In this study, by incorporating HECs into cerebral organoids derived from the same iPSC lines, we successfully produced cerebral organoids with induced microglia (iMG). Given their expression of CD309 and CD144, iPSC-derived HECs are closely related to erythroid-myeloid progenitors, from which microglia originate [45]. During the first month, the organoid medium was supplemented with factors necessary for macrophage/microglia generation (GM-CSF, IL-34). This supplementation was later discontinued after observing that CCOs intrinsically produced IL-34 and CX3CL1/Fractalkine, both of which are involved in microglia homeostasis.

The brain environment significantly influences the expression profile of microglial cells, distinguishing them from macrophages [17]. In our CCO model, iMG cells expressed key surface markers such as CX3CR1, P2RY12 and TMEM119, and displayed specific gene expression patterns consistent with previous studies [33, 39]. Notably, in our FACS experiments, iMG cells represented 0.42% [0.09–1.40] of the total cell population within CCOs, aligning with the levels of microglia observed in the adult human brain, which range from 0.5% to 16.6%, depending on the brain region [32]. While the iMG cells in our CCOs exhibited a rudimentary ramified morphology, they appeared to be functional. We demonstrated that these MG-like cells exhibited characteristics of natural microglia, including their ability to engulf neuronal synapses (PSD95), in agreement with previous findings [33, 54].

Thus, the model developed here offers an unprecedented opportunity to study functional human microglia phenotypes in both health and disease. We believe that the presence of vascular structures will further enable the integration of CCOs into microfluidic chips for advanced research applications. After validating our CCO model, we used it to explore the tumor microenvironment (TME) of GBM by coculturing CCOs with GSC lines. The TME includes various classes of non-neoplastic tissue-resident and infiltrating cells [36]. The non-neoplastic brain environment promotes electrochemical coupling between tumors and neurons, as well as metabolic interactions with astrocytes, which drive tumor growth [36]. Neurogenic niches have been proposed as a favorable environment for glioma cells [41], although the proximity of glioma cells to these niches may also reflect their stem cell origin [24]. In our observations, we did not find any preferential location or development of GSCs near the neurogenic niches of CCOs, particularly neural rosettes. Instead, we observed that in cocultures with CCOs, GSCs were frequently surrounded by numerous iMG cells (Iba1+), especially in the GBM-derived line (TG16), compared to the astrocytoma-derived line (TG20).

TAMs represent the largest population of non-cancerous cells in the GBM tumor microenvironment (TME), comprising up to 30–50% of the tumor mass [18]. This population includes a mix of resident microglia and macrophages derived from infiltrating monocytes [35]. While the infiltration of blood-derived TAMs can be inhibited using CSF1R inhibitors or chemokine receptor antagonists [42], resident microglia persist in the brain and can acquire TAM-like features. GBM cells can hijack microglial gene expression to support tumor growth [28], causing TAMs to adopt immunosuppressive properties [7]. Interactions between cancer cells and immune cells are known to drive the transition to mesenchymal-like states in GBM [19].

Our model allows for the specific investigation of how GSCs influence microglial cells. Indeed, we demonstrated that coculturing CCOs with GSCs alters the phenotype of iMG, particularly inducing them to adopt an immunosuppressive state, as evidenced by the increased expression of markers such as CD163, CD206, MSR1, and the TAM-associated gene SPP1. The higher expression levels of factors that influence TAMs in the TG16 line, such as CSF1 [42], CCL2 and POSTN [55], likely explain the more pronounced alterations in iMG in the presence of TG16 compared to TG20.

Additionally, GBM cells were often found in close proximity to vascular structures, suggesting vessel co-option in TG20 cells, while TG16 cells exhibited a tortuous morphology, characteristic of tumor-driven vascularization [22]. Moreover, endothelial cells in the cocultures showed elevated CD31 expression, a feature that has been previously reported in the tumor core [53].

Radiotherapy is administered to nearly all GBM patients, yet it may enhance the invasiveness of GSCs [15], which are highly resistant to radiation [6]. GSCs can infiltrate brain parenchyma and persist after surgical resection and radiotherapy [31]. Those located at the tumor margin are often exposed to lower doses of radiation than the central tumor mass. We demonstrated that GSCs proliferate in cocultures with conventional COs, and radiation exposure does not hinder their growth, consistent with previous studies using even higher doses of radiation [27]. Remarkably, we observed that irradiation of cocultures with CCOs actually promoted GSC proliferation. Although we cannot definitively pinpoint the specific cell population(s) responsible for this radiation-induced effect on GSCs, the enhanced recovery of GSCs was evident only in immunocompetent vascularized CCOs, not in conventional COs, suggesting a possible role for TAMs and/or vascular cells. This observation is reminiscent of the recurrences seen within 1–2 cm of the primary tumor border in GBM patients [11]. Similarly, in experimental GBM models in mice, initial depletion of tumor cells post-irradiation is often followed by significant recurrence, which is correlated with TME reorganization, particularly involving TAMs [1, 51]. Interactions between cancer and immune cells drive the transition to mesenchymal-like states in GBM [19].

Radiation also induces persistent reprogramming of microglia into a primed state [48] and expands TAM populations with an immunosuppressive profile [49]. Notably, we observed that iMGs were already reprogrammed toward an immunosuppressive TAM state in the presence of TG16 GSCs. A gene expression signature, known as the natural evolution signature (NES), has been identified in patients and is significantly associated with tumor evolution and macrophage polarization [52]. Intriguingly, several NES genes were upregulated after irradiation in TG16 GSCs cocultured with CCOs. Among these, POSTN plays a dual role in maintaining glioblastoma stem cells and promoting the immunosuppressive phenotype of microglia in GBM [55]. The immunosuppressive environment created by TAMs is responsible for the failure of immunotherapies [3]. Therefore, our model will be invaluable for elucidating the mechanisms driving recurrence after radiotherapy and for discovering new therapeutic strategies, particularly in the context of precision medicine.

In conclusion, our vascularized and immunocompetent CCO model offers a robust platform for investigating cellular interactions within the TME and assessing therapeutic responses. Additionally, it holds significant potential for advancing our understanding of human cerebral development and the disruptions that occur in diseases involving the endothelium and microglia.

## Acknowledgements

We are indebted to M.T. Garcia-Mitjavila, L. Morizur and A. Perrier for IPSC lines, advices in IPSC cultures and organoid generation, to V. Barroca and S. Devanand for mice experiments, to S. Messiaen for multiplex experiments, to F. Pflumio for giving immunodeficient mice and to V. Ménard for irradiation. JR is a fellow from CEA « CFR blanc ». JR is an ImmunoTools Award winner. This work was supported by La Ligue Contre le Cancer (comité des Hauts-de Seine), Les Entreprises Contre le Cancer (GEFLUC), Electricité de France, GIS FC3R (project No. 22FC3R-024) from funds managed by INSERM. IStem/CECS is supported by Association Française contre les Myopathies (AFM-Téléthon).

## References

1. 1. Akkari L, Bowman RL, Tessier J, Klemm F, Handgraaf SM, de Groot M, Quail DF, Tillard L, Gadiot J, Huse JT, Brandsma D, Westerga J, Watts C, Joyce JA (2020) Dynamic changes in glioma macrophage populations after radiotherapy reveal CSF-1R inhibition as a strategy to overcome resistance. Sci Transl Med 12:eaaw7843. doi: 10.1126/scitranslmed.aaw7843

2. Amankulor NM, Kim Y, Arora S, Kargl J, Szulzewsky F, Hanke M, Margineantu DH, Rao A, Bolouri H, Delrow J, Hockenbery D, Houghton AM, Holland EC (2017) Mutant IDH1 regulates the tumor-associated immune system in gliomas. Genes Dev 31:774–786. doi: 10.1101/gad.294991.116

3. Antonucci L, Canciani G, Mastronuzzi A, Carai A, Del Baldo G, Del Bufalo F (2022) CAR-T Therapy for Pediatric High-Grade Gliomas: Peculiarities, Current Investigations and Future Strategies. Front Immunol 13:867154. doi: 10.3389/fimmu.2022.867154

4. Atkins MH, Scarfò R, McGrath KE, Yang D, Palis J, Ditadi A, Keller GM (2021) Modeling human yolk sac hematopoiesis with pluripotent stem cells. Journal of Experimental Medicine 219:e20211924. doi: 10.1084/jem.20211924

5. Azzoni E, Conti V, Campana L, Dellavalle A, Adams RH, Cossu G, Brunelli S (2014) Hemogenic endothelium generates mesoangioblasts that contribute to several mesodermal lineages in vivo. Development 141:1821–1834. doi: 10.1242/dev.103242

6. Bao S, Wu Q, McLendon RE, Hao Y, Shi Q, Hjelmeland AB, Dewhirst MW, Bigner DD, Rich JN (2006) Glioma stem cells promote radioresistance by preferential activation of the DNA damage response. Nature 444:756–60. doi: 10.1038/nature05236

7. Blériot C, Dunsmore G, Alonso-Curbelo D, Ginhoux F (2024) A temporal perspective for tumor-associated macrophage identities and functions. Cancer Cell 42:747–758. doi: 10.1016/j.ccell.2024.04.002

8. Cakir B, Xiang Y, Tanaka Y, Kural MH, Parent M, Kang Y-J, Chapeton K, Patterson B, Yuan Y, He C-S, Raredon MSB, Dengelegi J, Kim K-Y, Sun P, Zhong M, Lee S, Patra P, Hyder F, Niklason LE, Lee S-H, Yoon Y-S, Park I-H (2019) Engineering of human brain organoids with a functional vascular-like system. Nat Methods 16:1169–1175. doi: 10.1038/s41592-019-0586-5

9. Carmeliet P, Tessier-Lavigne M (2005) Common mechanisms of nerve and blood vessel wiring. Nature 436:193–200. doi: 10.1038/nature03875

10. Chiaradia I, Lancaster MA (2020) Brain organoids for the study of human neurobiology at the interface of in vitro and in vivo. Nat Neurosci 23:1496–1508. doi: 10.1038/s41593-020-00730-3

11. Cuddapah VA, Robel S, Watkins S, Sontheimer H (2014) A neurocentric perspective on glioma invasion. Nat Rev Neurosci 15:455–465. doi: 10.1038/nrn3765

12. Daneman R, Prat A (2015) The Blood–Brain Barrier. Cold Spring Harb Perspect Biol 7:a020412. doi: 10.1101/cshperspect.a020412

13. Dao L, You Z, Lu L, Xu T, Sarkar AK, Zhu H, Liu M, Calandrelli R, Yoshida G, Lin P, Miao Y, Mierke S, Kalva S, Zhu H, Gu M, Vadivelu S, Zhong S, Huang LF, Guo Z (2024) Modeling blood-brain barrier formation and cerebral cavernous malformations in human PSC-derived organoids. Cell Stem Cell 31:818–833.e11. doi: 10.1016/j.stem.2024.04.019

14. El-Habr EA, Dubois LG, Burel-Vandenbos F, Bogeas A, Lipecka J, Turchi L, Lejeune F-X, Coehlo PLC, Yamaki T, Wittmann BM, Fareh M, Mahfoudhi E, Janin M, Narayanan A, Morvan-Dubois G, Schmitt C, Verreault M, Oliver L, Sharif A, Pallud J, Devaux B, Puget S, Korkolopoulou P, Varlet P, Ottolenghi C, Plo I, Moura-Neto V, Virolle T, Chneiweiss H, Junier M-P (2017) A driver role for GABA metabolism in controlling stem and proliferative cell state through GHB production in glioma. Acta Neuropathol 133:645–660. doi: 10.1007/s00401-016-1659-5

15. Gauthier LR, Saati M, Bensalah-Pigeon H, Ben M’Barek K, Gitton-Quent O, Bertrand R, Busso D, Mouthon M-A, Collura A, Junier M-P, Chneiweiss H, Pineda JR, Boussin FD (2020) The HIF1α/JMY pathway promotes glioblastoma stem-like cell invasiveness after irradiation. Sci Rep 10:18742. doi: 10.1038/s41598-020-75300-5

16. Ginhoux F, Greter M, Leboeuf M, Nandi S, See P, Gokhan S, Mehler MF, Conway SJ, Ng LG, Stanley ER, Samokhvalov IM, Merad M (2010) Fate mapping analysis reveals that adult microglia derive from primitive macrophages. Science 330:841–845. doi: 10.1126/science.1194637

17. Gosselin D, Skola D, Coufal NG, Holtman IR, Schlachetzki JCM, Sajti E, Jaeger BN, O’Connor C, Fitzpatrick C, Pasillas MP, Pena M, Adair A, Gonda DD, Levy ML, Ransohoff RM, Gage FH, Glass CK (2017) An environment-dependent transcriptional network specifies human microglia identity. Science 356:eaal3222. doi: 10.1126/science.aal3222

18. Gutmann DH, Kettenmann H (2019) Microglia/Brain Macrophages as Central Drivers of Brain Tumor Pathobiology. Neuron 104:442–449. doi: 10.1016/j.neuron.2019.08.028

19. Hara T, Chanoch-Myers R, Mathewson ND, Myskiw C, Atta L, Bussema L, Eichhorn SW, Greenwald AC, Kinker GS, Rodman C, Gonzalez Castro LN, Wakimoto H, Rozenblatt-Rosen O, Zhuang X, Fan J, Hunter T, Verma IM, Wucherpfennig KW, Regev A, Suvà ML, Tirosh I (2021) Interactions between cancer cells and immune cells drive transitions to mesenchymal-like states in glioblastoma. Cancer Cell 39:779–792.e11. doi: 10.1016/j.ccell.2021.05.002

20. Jeitany M, Pineda JR, Liu Q, Porreca RM, Hoffschir F, Desmaze C, Silvestre DC, Mailliet P, Junier MP, Londono-Vallejo A, Segal-Bendirdjian E, Chneiweiss H, Boussin FD (2015) A preclinical mouse model of glioma with an alternative mechanism of telomere maintenance (ALT). Int J Cancer 136:1546–58. doi: 10.1002/ijc.29171

21. Jeon H-M, Kim J-Y, Cho HJ, Lee WJ, Nguyen D, Kim SS, Oh YT, Kim H-J, Jung C-W, Pinero G, Joshi T, Hambardzumyan D, Sakaguchi T, Hubert CG, McIntyre TM, Fine HA, Gladson CL, Wang B, Purow BW, Park JB, Park MJ, Nam D-H, Lee J (2023) Tissue factor is a critical regulator of radiation therapy-induced glioblastoma remodeling. Cancer Cell 41:1480–1497.e9. doi: 10.1016/j.ccell.2023.06.007

22. Jhaveri N, Chen TC, Hofman FM (2016) Tumor vasculature and glioma stem cells: Contributions to glioma progression. Cancer Lett 380:545–551. doi: 10.1016/j.canlet.2014.12.028

23. Krieger TG, Tirier SM, Park J, Jechow K, Eisemann T, Peterziel H, Angel P, Eils R, Conrad C (2020) Modeling glioblastoma invasion using human brain organoids and single-cell transcriptomics. Neuro Oncol 22:1138–1149. doi: 10.1093/neuonc/noaa091

24. Kusne Y, Sanai N (2015) The SVZ and Its Relationship to Stem Cell Based Neuro-oncogenesis. Adv Exp Med Biol 853:23–32. doi: 10.1007/978-3-319-16537-0_2

25. Lancaster MA, Renner M, Martin C-A, Wenzel D, Bicknell LS, Hurles ME, Homfray T, Penninger JM, Jackson AP, Knoblich JA (2013) Cerebral organoids model human brain development and microcephaly. Nature 501:373–379. doi: 10.1038/nature12517

26. Li W, Germain RN, Gerner MY (2019) High-dimensional cell-level analysis of tissues with Ce3D multiplex volume imaging. Nature Protocols 14:1708–1733. doi: 10.1038/s41596-019-0156-4

27. Linkous A, Balamatsias D, Snuderl M, Edwards L, Miyaguchi K, Milner T, Reich B, Cohen-Gould L, Storaska A, Nakayama Y, Schenkein E, Singhania R, Cirigliano S, Magdeldin T, Lin Y, Nanjangud G, Chadalavada K, Pisapia D, Liston C, Fine HA (2019) Modeling Patient-Derived Glioblastoma with Cerebral Organoids. Cell Rep 26:3203–3211.e5. doi: 10.1016/j.celrep.2019.02.063

28. 28. Maas SLN, Abels ER, Van De Haar LL, Zhang X, Morsett L, Sil S, Guedes J, Sen P, Prabhakar S, Hickman SE, Lai CP, Ting DT, Breakefield XO, Broekman MLD, El Khoury J (2020) Glioblastoma hijacks microglial gene expression to support tumor growth. J Neuroinflammation 17:120. doi: 10.1186/s12974-020-01797-2

29. Mansour AA, Gonçalves JT, Bloyd CW, Li H, Fernandes S, Quang D, Johnston S, Parylak SL, Jin X, Gage FH (2018) An in vivo model of functional and vascularized human brain organoids. Nat Biotechnol 36:432–441. doi: 10.1038/nbt.4127

30. Marcelo KL, Goldie LC, Hirschi KK (2013) Regulation of endothelial cell differentiation and specification. Circ Res 112:1272–1287. doi: 10.1161/CIRCRESAHA.113.300506

31. Marhuenda E, Fabre C, Zhang C, Martin-Fernandez M, Iskratsch T, Saleh A, Bauchet L, Cambedouzou J, Hugnot J-P, Duffau H, Dennis JW, Cornu D, Bakalara N (2021) Glioma stem cells invasive phenotype at optimal stiffness is driven by MGAT5 dependent mechanosensing. J Exp Clin Cancer Res 40:139. doi: 10.1186/s13046-021-01925-7

32. Mittelbronn M, Dietz K, Schluesener HJ, Meyermann R (2001) Local distribution of microglia in the normal adult human central nervous system differs by up to one order of magnitude. Acta Neuropathol 101:249–255. doi: 10.1007/s004010000284

33. Park DS, Kozaki T, Tiwari SK, Moreira M, Khalilnezhad A, Torta F, Olivié N, Thiam CH, Liani O, Silvin A, Phoo WW, Gao L, Triebl A, Tham WK, Gonçalves L, Kong WT, Raman S, Zhang XM, Dunsmore G, Dutertre CA, Lee S, Ong JM, Balachander A, Khalilnezhad S, Lum J, Duan K, Lim ZM, Tan L, Low I, Utami KH, Yeo XY, Di Tommaso S, Dupuy J-W, Varga B, Karadottir RT, Madathummal MC, Bonne I, Malleret B, Binte ZY, Wei Da N, Tan Y, Wong WJ, Zhang J, Chen J, Sobota RM, Howland SW, Ng LG, Saltel F, Castel D, Grill J, Minard V, Albani S, Chan JKY, Thion MS, Jung SY, Wenk MR, Pouladi MA, Pasqualini C, Angeli V, Cexus ONF, Ginhoux F (2023) iPS-cell-derived microglia promote brain organoid maturation via cholesterol transfer. Nature 623:397–405. doi: 10.1038/s41586-023-06713-1

34. Paşca AM, Sloan SA, Clarke LE, Tian Y, Makinson CD, Huber N, Kim CH, Park J-Y, O’Rourke NA, Nguyen KD, Smith SJ, Huguenard JR, Geschwind DH, Barres BA, Paşca SP (2015) Functional cortical neurons and astrocytes from human pluripotent stem cells in 3D culture. Nat Methods 12:671–678. doi: 10.1038/nmeth.3415

35. Pombo Antunes AR, Scheyltjens I, Lodi F, Messiaen J, Antoranz A, Duerinck J, Kancheva D, Martens L, De Vlaminck K, Van Hove H, Kjølner Hansen SS, Bosisio FM, Van der Borght K, De Vleeschouwer S, Sciot R, Bouwens L, Verfaillie M, Vandamme N, Vandenbroucke RE, De Wever O, Saeys Y, Guilliams M, Gysemans C, Neyns B, De Smet F, Lambrechts D, Van Ginderachter JA, Movahedi K (2021) Single-cell profiling of myeloid cells in glioblastoma across species and disease stage reveals macrophage competition and specialization. Nat Neurosci 24:595–610. doi: 10.1038/s41593-020-00789-y

36. Read RD, Tapp ZM, Rajappa P, Hambardzumyan D (2024) Glioblastoma microenvironment-from biology to therapy. Genes Dev. doi: 10.1101/gad.351427.123

37. Renner M, Lancaster MA, Bian S, Choi H, Ku T, Peer A, Chung K, Knoblich JA (2017) Self-organized developmental patterning and differentiation in cerebral organoids. EMBO J 36:1316–1329. doi: 10.15252/embj.201694700

38. Rosińska S, Gavard J (2021) Tumor Vessels Fuel the Fire in Glioblastoma. Int J Mol Sci 22:6514. doi: 10.3390/ijms22126514

39. Schafer ST, Mansour AA, Schlachetzki JCM, Pena M, Ghassemzadeh S, Mitchell L, Mar A, Quang D, Stumpf S, Ortiz IS, Lana AJ, Baek C, Zaghal R, Glass CK, Nimmerjahn A, Gage FH (2023) An in vivo neuroimmune organoid model to study human microglia phenotypes. Cell 186:2111–2126.e20. doi: 10.1016/j.cell.2023.04.022

40. Silvestre DC, Pineda JR, Hoffschir F, Studler JM, Mouthon MA, Pflumio F, Junier MP, Chneiweiss H, Boussin FD (2011) Alternative lengthening of telomeres in human glioma stem cells. Stem Cells 29:440–51

41. Sinnaeve J, Mobley BC, Ihrie RA (2018) Space Invaders: Brain Tumor Exploitation of the Stem Cell Niche. Am J Pathol 188:29–38. doi: 10.1016/j.ajpath.2017.08.029

42. Stafford JH, Hirai T, Deng L, Chernikova SB, Urata K, West BL, Brown JM (2016) Colony stimulating factor 1 receptor inhibition delays recurrence of glioblastoma after radiation by altering myeloid cell recruitment and polarization. Neuro Oncol 18:797–806. doi: 10.1093/neuonc/nov272

43. Sun X-Y, Ju X-C, Li Y, Zeng P-M, Wu J, Zhou Y-Y, Shen L-B, Dong J, Chen Y-J, Luo Z-G (2022) Generation of vascularized brain organoids to study neurovascular interactions. Elife 11:e76707. doi: 10.7554/eLife.76707

44. Thion MS, Ginhoux F, Garel S (2018) Microglia and early brain development: An intimate journey. Science 362:185–189. doi: 10.1126/science.aat0474

45. 45. Vargas-Valderrama A, Ponsen A-C, Le Gall M, Clay D, Jacques S, Manoliu T, Rouffiac V, Ser-le-Roux K, Quivoron C, Louache F, Uzan G, Mitjavila-Garcia M-T, Oberlin E, Guenou H (2022) Endothelial and hematopoietic hPSCs differentiation via a hematoendothelial progenitor. Stem Cell Res Ther 13:254. doi: 10.1186/s13287-022-02925-w

46. Velasco S, Kedaigle AJ, Simmons SK, Nash A, Rocha M, Quadrato G, Paulsen B, Nguyen L, Adiconis X, Regev A, Levin JZ, Arlotta P (2019) Individual brain organoids reproducibly form cell diversity of the human cerebral cortex. Nature 570:523–527. doi: 10.1038/s41586-019-1289-x

47. Vollmann-Zwerenz A, Leidgens V, Feliciello G, Klein CA, Hau P (2020) Tumor Cell Invasion in Glioblastoma. Int J Mol Sci 21. doi: 10.3390/ijms21061932

48. 48. Voshart DC, Oshima T, Jiang Y, van der Linden GP, Ainslie AP, Reali Nazario L, van Buuren-Broek F, Scholma AC, van Weering HRJ, Brouwer N, Sewdihal J, Brouwer U, Coppes RP, Holtman IR, Eggen BJL, Kooistra SM, Barazzuol L (2024) Radiotherapy induces persistent innate immune reprogramming of microglia into a primed state. Cell Rep 43:113764. doi: 10.1016/j.celrep.2024.113764

49. Wang L, Dou X, Chen S, Yu X, Huang X, Zhang L, Chen Y, Wang J, Yang K, Bugno J, Pitroda S, Ding X, Piffko A, Si W, Chen C, Jiang H, Zhou B, Chmura SJ, Luo C, Liang HL, He C, Weichselbaum RR (2023) YTHDF2 inhibition potentiates radiotherapy antitumor efficacy. Cancer Cell 41:1294–1308.e8. doi: 10.1016/j.ccell.2023.04.019

50. Wang W, Li T, Cheng Y, Li F, Qi S, Mao M, Wu J, Liu Q, Zhang X, Li X, Zhang L, Qi H, Yang L, Yang K, He Z, Ding S, Qin Z, Yang Y, Yang X, Luo C, Guo Y, Wang C, Liu X, Zhou L, Liu Y, Kong W, Miao J, Ye S, Luo M, An L, Wang L, Che L, Niu Q, Ma Q, Zhang X, Zhang Z, Hu R, Feng H, Ping Y-F, Bian X-W, Shi Y (2024) Identification of hypoxic macrophages in glioblastoma with therapeutic potential for vasculature normalization. Cancer Cell 42:815–832.e12. doi: 10.1016/j.ccell.2024.03.013

51. 51. Watson SS, Duc B, Kang Z, de Tonnac A, Eling N, Font L, Whitmarsh T, Massara M, iMAXT Consortium, Bodenmiller B, Hausser J, Joyce JA (2024) Microenvironmental reorganization in brain tumors following radiotherapy and recurrence revealed by hyperplexed immunofluorescence imaging. Nat Commun 15:3226. doi: 10.1038/s41467-024-47185-9

52. Wu L, Wu W, Zhang J, Zhao Z, Li L, Zhu M, Wu M, Wu F, Zhou F, Du Y, Chai R-C, Zhang W, Qiu X, Liu Q, Wang Z, Li J, Li K, Chen A, Jiang Y, Xiao X, Zou H, Srivastava R, Zhang T, Cai Y, Liang Y, Huang B, Zhang R, Lin F, Hu L, Wang X, Qian X, Lv S, Hu B, Zheng S, Hu Z, Shen H, You Y, Verhaak RGW, Jiang T, Wang Q (2022) Natural Coevolution of Tumor and Immunoenvironment in Glioblastoma. Cancer Discov 12:2820–2837. doi: 10.1158/2159-8290.CD-22-0196

53. Xie Y, He L, Lugano R, Zhang Y, Cao H, He Q, Chao M, Liu B, Cao Q, Wang J, Jiao Y, Hu Y, Han L, Zhang Y, Huang H, Uhrbom L, Betsholtz C, Wang L, Dimberg A, Zhang L (2021) Key molecular alterations in endothelial cells in human glioblastoma uncovered through single-cell RNA sequencing. JCI Insight 6:e150861. doi: 10.1172/jci.insight.150861

54. Zhang W, Jiang J, Xu Z, Yan H, Tang B, Liu C, Chen C, Meng Q (2023) Microglia-containing human brain organoids for the study of brain development and pathology. Mol Psychiatry 28:96–107. doi: 10.1038/s41380-022-01892-1

55. Zhou W, Ke SQ, Huang Z, Flavahan W, Fang X, Paul J, Wu L, Sloan AE, McLendon RE, Li X, Rich JN, Bao S (2015) Periostin secreted by glioblastoma stem cells recruits M2 tumour-associated macrophages and promotes malignant growth. Nat Cell Biol 17:170–182. doi: 10.1038/ncb3090

